# Piezo1 Mediates Keratinocyte Mechanotransduction

**DOI:** 10.1101/2020.07.19.211086

**Authors:** Francie Moehring, Alexander R. Mikesell, Katelyn E. Sadler, Anthony D. Menzel, Cheryl L. Stucky

## Abstract

Epidermal keratinocytes mediate touch sensation by detecting and encoding tactile information to sensory neurons. However, the specific mechanotransducers that enable keratinocytes to respond to mechanical stimulation are unknown. Here, we found that the mechanically-gated ion channel Piezo1 is the major keratinocyte mechanotransducer. Keratinocyte expression of Piezo1 is critical for normal sensory afferent firing and behavioral responses to mechanical stimuli.

## Main

In the last decade, sensory biologists have demonstrated a requirement for non-neuronal cells in normal touch sensation^1–5^. Work from our lab and others has revealed that keratinocytes, which constitute > 90% of epidermal cells^6^, are inherently responsive to mechanical force^3^, modulate sensory afferent firing in response to mechanical stimuli^7^, and are required for normal touch sensation^3^. The ability of keratinocytes to respond to force indicates that they must possess one or more mechanically sensitive channels, but the specific keratinocyte mechanotransducers have not yet been identified. Piezo1 and Piezo2 are mechanically-gated, non-selective cation channels^8^; while Piezo2 is required for mechanotransduction in sensory neurons and Merkel cells^4,9^, a definitive role for Piezo1 in mechanosensation has never been identified. Because Piezo1 is highly expressed in mouse skin^8^, we hypothesized that this channel is the primary mechanotransducer in keratinocytes.

To test this hypothesis, we generated epidermal cell-specific Piezo1 knockout mice by crossing Keratin14- (K14) Cre and Piezo1^flox/flox^ mice^10^. Successful knockout of the channel was verified with *in vitro* calcium imaging of primary epidermal keratinocytes; keratinocytes isolated from wildtype controls responded robustly to the Piezo1-specific chemical agonist Yoda1^11^ while keratinocytes isolated from Piezo1 knockout mice (Piezo1cKO) were virtually unresponsive (**Fig 1A-C**). To determine if direct activation of epidermal Piezo1 is sufficient to induce *in vivo* behavioral responses, we injected Yoda1 into the hindpaw of wildtype and Piezo1cKO mice. Yoda1 induced dose dependent paw attending responses in wildtype mice (**Fig 1D**) but had no effect in Piezo1cKO mice (**Fig 1E**), suggesting that the observed attending behaviors were dependent on epidermally-expressed Piezo1. We next determined whether epidermal Piezo1 is required for normal behavioral tactile sensation. Piezo1cKO mice displayed muted behavioral responses to innocuous punctate (**Fig 1F-H**), innocuous dynamic (**Fig 1I**), and noxious punctate (**Fig 1J**) stimulation. Piezo1cKO mice did not exhibit general somatosensory deficits as the mutants were just as sensitive as wildtype controls to heat (**Fig 1K**) and cold (**Fig 1L**) stimulation. Likewise, these behavioral changes did not result from a change in epidermal morphology, as the stratum corneum and stratum spinosum were indistinguishable between Piezo1cKO and wildtype mice (**Fig 1M-N**). These data indicate that epidermal Piezo1 is critical for normal innocuous and noxious mechanosensation.

**Figure 1.**
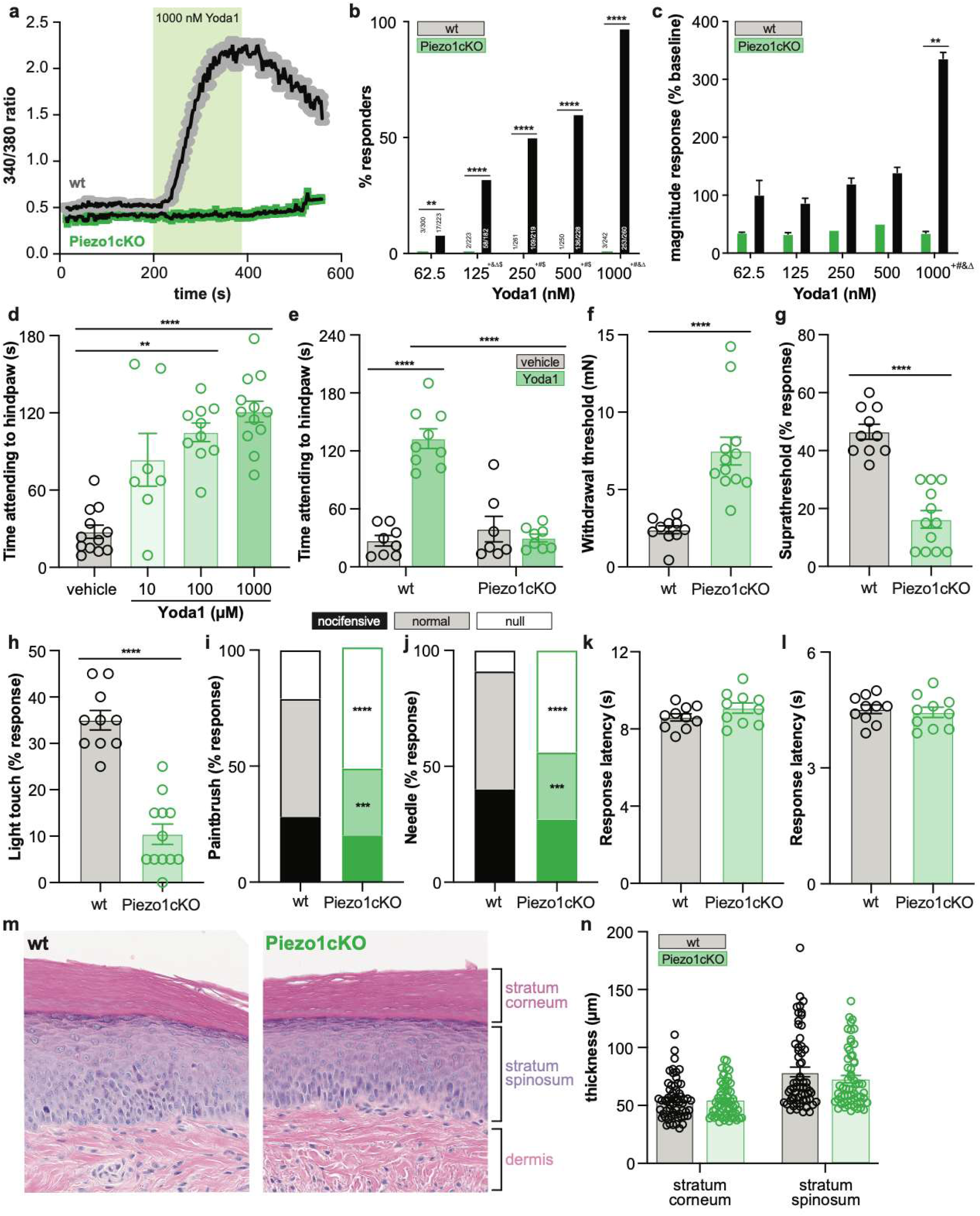
Epidermal Piezo1 is required for Yoda1 induced behavioral responses and for normal innocuous and noxious mechanosensation. **a**. Calcium flux in wildtype (wt) and Piezo1cKO keratinocytes in response to 1,000 nM Yoda1. **b**. Percentage of wildtype and Piezo1cKO keratinocytes that respond to extracellular Yoda1; cells from n=3 mice; bars are group averages. **c**. Peak calcium responses to extracellular Yoda1. **d**. Yoda1-induced attending behaviors in wildtype mice. **e**. Yoda1 (1 mM) induced attending behaviors in wildtype and Piezo1cKO mice. **f**. von Frey mechanical thresholds of wildtype and Piezo1cKO mice. **g**. Wildtype and Piezo1cKO responses to repeated suprathreshold (3.61 mN) von Frey filament stimulation. **h**. Wildtype and Piezo1cKO responses to repeated static light touch (0.6 mN von Frey filament) stimulation. **i**. Response characterization to paintbrush swiping across hindpaw; n=10-12; bars are group averages **j**. Response characterization to noxious needle hindpaw stimulation; n=10-12; bars are group averages. **k**. Withdrawal latency to radiant heat hindpaw stimulation. **l**. Withdrawal latency to dry ice hindpaw stimulation. **m**. H&E stained skin from the hindpaw of wildtype and Piezo1cKO mice. **n**. Quantification of individual epidermal layer thickness. All data are mean ± SEM unless otherwise stated. Post-hoc comparisons for all panels: **P<0.01, ***P<0.001, ****P<0.0001.

Since epidermal deletion of Piezo1 reduced behavioral responses to mechanical stimulation, we next used *ex vivo* tibial nerve recordings to determine whether epidermal Piezo1 deletion affects mechanically evoked sensory afferent firing. Slowly adapting (SA) Aβ fibers from Piezo1cKO mice fired fewer action potentials during receptive field mechanical simulation than fibers isolated from wildtype control mice (**Fig 2A, D, and G**). Decreases in mechanically-induced firing were specifically observed during the sustained phase of stimulus application (**Fig 2G**). No difference in mechanical thresholds were noted between Piezo1cKO and wildtype Aβ fibers (**Fig 2J and K**). Aδ fibers from Piezo1cKO mice also fired fewer action potentials than fibers from wildtype mice during mechanical stimulation (**Fig 2B**). However, differences in firing were not restricted to the sustained phase of stimulus application but also occurred at the onset of the stimulus (**Fig 2H**). Corresponding with this, the mechanical thresholds of Piezo1cKO Aδ fibers were significantly higher than wildtype fibers (**Fig 2J and I**). No differences in C fiber firing frequency or mechanical thresholds were observed between Piezo1cKO and wildtype control preparations (**Fig 2C, F, I, J, and M)**. These data demonstrate that normal mechanically induced firing of myelinated cutaneous SA-Aβ and Aδ primary afferents depends on the epidermal expression of Piezo1. Differences in firing frequency were most pronounced at higher forces, likely reflecting the summation of keratinocyte ATP signaling at the sensory afferent terminals^3^. Because we utilized a K14 promoter to target Piezo1 deletion, we cannot exclude the contribution of K14 expressing Merkel cells in our experiments^1^. However, we believe that our results are relatively specific to keratinocytes for two reasons: (1) Merkel cells express minimal Piezo1^1,12^, and (2) Merkel cell mechanical sensitivity is exclusively mediated by Piezo2; Piezo2 deletion renders Merkel cells mechanically insensitive ^4^. We were surprised to observe no effect of epidermal Piezo1 deletion on C fiber mechanical firing given that optogenetic keratinocyte inhibition decreases C fiber mechanical firing ^7^. Because many C fibers terminate in more superficial layers of the epidermis than Aδ and Aβ fibers, it is possible that differentiated keratinocytes of the outer epidermis rely on a different mechanotransducer to encode tactile information to C fibers^13,14^.

**Figure 2.**
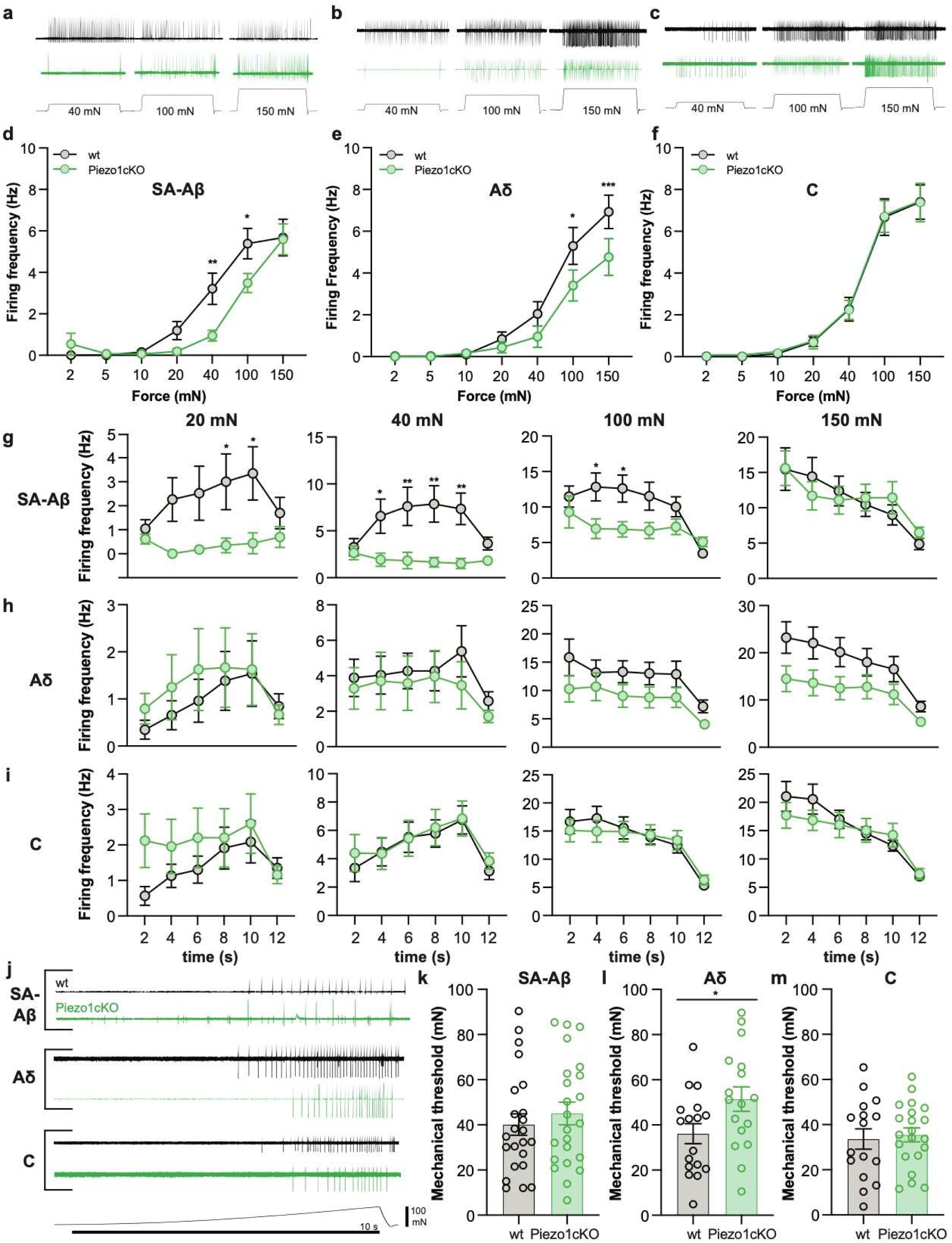
Normal mechanically-induced primary afferent firing requires epidermal Piezo1 expression. *Ex vivo* tibial nerve recordings of Piezo1cKO and wildtype (wt) **a**. SA-Aβ fibers, **b**. Aδ fibers, and **c**. C fibers. Mean mechanically-induced firing rates of **d**. SA-Aβ fibers, **e**. Aδ fibers, and **f**. C fibers. Binned firing rates during sustained force applications to receptive field of **g**. SA-Aβ fibers, **h**. Aδ fibers, and **i**. C fibers. **j**. Single unit firing in response to mechanical force ramp (0 to 100 mN; 10 s). Mechanical thresholds of **k**. SA-Aβ fibers, **l**. Aδ fibers, and **m**. C fibers. All data are mean ± SEM. Post-hoc comparisons for all panels: *P<0.05, **P<0.01, ***P<0.001; fibers from n=13-16 mice.

We concluded by investigating if Piezo1 deletion specifically decreased keratinocyte mechanical sensitivity. In whole-cell current clamp mode, mechanical probing of wildtype keratinocytes led to force-dependent membrane depolarization, a response that was decreased in Piezo1cKO keratinocytes (**Fig 3A**). Recordings in whole cell voltage-clamp mode revealed that wildtype keratinocytes respond to focal mechanical stimulation with graded inward currents (**Fig 3B and 4C**). The majority of wildtype keratinocytes exhibited rapidly-adapting currents upon mechanical stimulation (**Fig 3D**). In contrast, a large number (> 65%) of Piezo1cKO keratinocytes were unresponsive to mechanical probing (i.e. mechanically insensitive), indicating that in these keratinocytes, Piezo1 is the primary mechanotransducer. The minority of Piezo1cKO keratinocytes that retained mechanical sensitivity exhibited smaller mechanically-evoked currents than wildtype controls (**Fig 3B-D**), but exhibited no difference in mechanical threshold, suggesting that Piezo1 is a secondary mechanotransducer in these cells, rather than the channel that sets the mechanical threshold (**Fig. 3E**). There was no difference in resting membrane capacitance or resting membrane potential between Piezo1cKO and wildtype keratinocytes, indicating that Piezo1 deletion does not disrupt resting electrochemical properties (**Fig. 3F and G**). These data indicate that Piezo1 is the major keratinocyte mechanotransducer. This study is the first to demonstrate that Piezo1 is required for normal touch sensation. Our results show that keratinocyte-expressed Piezo1 enables these cells to encode a wide range of intensities and qualities of mechanical stimuli to primary afferents, and thus, critically mediates tactile behavioral responses. Future studies should examine how Piezo1 sensitization and resulting keratinocyte activity contribute to mechanical allodynia and hyperalgesia in peripheral tissue injury.

**Figure 3.**
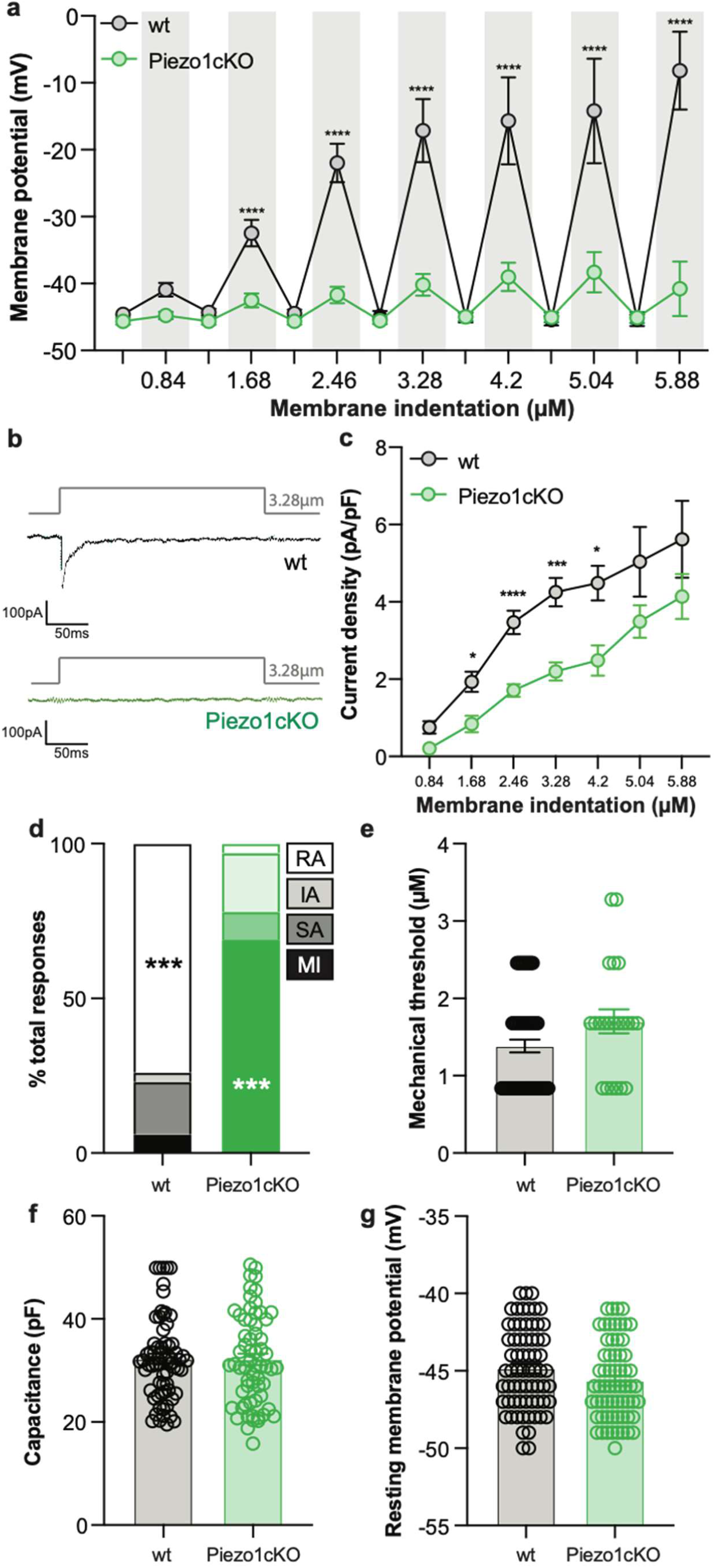
Piezo1cKO keratinocytes exhibit a decrease in mechanically-evoked inward currents and an increase in mechanically insensitive cells. **a**. Mechanically-induced membrane depolarization in isolated Piezo1cKO and wildtype (wt) keratinocytes. **b**. Whole cell mechanically-induced currents. **c**. Mean mechanical current densities. **d**. Characterization of keratinocyte mechanical currents as rapidly adapting (RA), intermediately adapting (IA), slowly adapting (SA), or mechanically insensitive (MI); bars are group averages **e**. Keratinocyte mechanical thresholds. **F**. Keratinocyte cell capacitance. **g**. Keratinocyte resting membrane potentials. All data are mean ± SEM unless otherwise stated. Post-hoc comparisons for all panels: *P<0.05, ***P<0.001, ****P<0.0001; cells from n=8-10 mice.

## Methods

### Animals

To target epidermal keratinocytes, a *Keratin14* (K14) Cre driver was used, as *Keratin14* is expressed in all keratinocytes as early as E9.5 ^15–17^. These mice were mated with Piezo1^fl/fl^ animals to produce offspring that lacked Piezo1 in K14 positive cells and were genotyped as either *K14Cre*^*+*^ *Piezo1*^*fl/fl*^ (Piezo1cKO) or *K14Cre*^*-*^ *Piezo1*^*fl/fl*^ (wildtype). For all studies a mixture of male and female mice aged 6 - 20 weeks were used. Male and female mice were analyzed separately, and no sex differences were observed. Therefore, data shown in graphs show combined results of both sexes.

Animals had *ad libitum* access to food and water and were housed in a climate-controlled room with a 12:12 light:dark cycle, on Sani-Chips® aspen wood chip bedding (P.J. Murphy Forest and Products, New Jersey) with a single pack of ENVIROPAK nesting material (W.F Fisher & Son, Inc, New Jersey). All animals were group housed with a minimum of 3 mice per cage. All animal procedures were strictly adhered according to the NIH Guide for the Care and Use of Laboratory animals and were performed in accordance with the Institutional Animal Care and Use Committee at the Medical College of Wisconsin (approval #0383). This manuscript adheres to the applicable ARRIVE guidelines.

### Primary Keratinocyte Cell culture

Primary keratinocytes were cultured from glabrous hindpaw tissue as previously described (Moehring et al., 2018). Briefly, isolated glabrous hindpaw skin was incubated at room temperature (RT) in 10 mg/mL dispase (Gibco, ThermoFisher Scientific, Waltham, MA) for 45 min. Following the dispase incubation, the epidermal sheet was separated from the dermis and incubated at RT in 50% EDTA (Sigma-Aldrich) and 0.05% trypsin (Sigma) in Hanks’ Balanced Salt Solution without calcium chloride, magnesium chloride and magnesium sulfate (Gibco) for 27 min. After 27 min, 15% heat inactivated fetal bovine serum (ThermoFisher Scientific, Carlsbad, CA) was added and the epidermal sheets were rubbed against the base of a petri dish to separate the keratinocytes. Keratinocytes were grown for 3 days in Epilife media (Gibco) supplemented with 1% human keratinocyte growth supplement (Gibco), 0.2% GibcoAmphotericin B (250 µg/mL of Amphotericin B and 205 µg/mL sodium deoxycholate, Gibco) and 0.25% penicillin-streptomycin (Gibco) on laminin coated coverslips. Plates were kept at 37°C and 5% CO2 conditions. Growth media was exchanged every 2 days.

### Calcium Imaging

Calcium imaging was performed on keratinocytes on their third day in culture. Keratinocytes were loaded with 2.5 µg/mL Fura-2-AM, a dual-wavelength ratiometric calcium indicator dye, in 2% BSA for 45 min at RT then washed with extracellular buffer for 30 min. Keratinocytes were superfused with RT extracellular buffer (pH 7.4, 320 Osm) containing (in mM) 150 NaCl, 10 HEPES, 8 glucose, 5.6 KCl, 2 CaCl2, and 1 MgCl2, and viewed on a Nikon Eclipse TE200 inverted microscope. Nikon elements software (Nikon Instruments, Melville, NY) was used to capture fluorescence images at 340 and 380 nm. Responsive cells were those that exhibited >30% increase in 340/380nm ratio from baseline. Yoda-1 was used at 62.5, 125, 250, 500 and 1,000 (nM) concentrations in extracellular buffer for the dose response curve. Yoda-1 was applied for 3 min and washed out for 3 min.

### Behavioral Assays

For all (spontaneous and evoked) behavior experiments the experimenter was blinded to genotype throughout testing and data entry. Animals were tested between 8am and 1pm and were allowed to acclimate for at least an hour to the new surroundings and experimenter before any behavior testing was performed.

#### Mechanical sensitivity

A battery of different mechanical assays using various mechanical stimuli were utilized to determine the mechanical sensitivity of the of Piezo1cKO and wildtype littermate controls. Using the Up-Down method and a series of calibrated von Frey filaments ranging from 0.38-37 mN, mechanical thresholds of the glabrous hindpaw skin were assessed^18,19^. Additionally, the hindpaw skin was stimulated 10 times using a 3.61 mN von Frey Filament in the suprathreshold assay, and using a 0.6 mN von Frey Filament in the static light touch assay. The number of stimulus-evoked paw withdrawals were recorded^20^. Furthermore, we utilized a paintbrush that was stroked 5 times across the hindpaw and responses were categorized as: normal/innocuous (simple withdrawal of the paw), noxious (elevating the paw for extended periods of time, flicking and licking of the paw), and null responses (no withdrawal)^21^. Lastly, noxious mechanical sensitivity was assessed using the needle assay^22,23^, where responses were categorized similar to the paintbrush assay.

To test <u>heat sensitivity</u>, mice were placed in a small plexiglass enclosures on top of a glass plate, and a focal radiant heat source was applied to the plantar hindpaw. The response latency to hind paw withdrawal from the heat stimulus was quantified^24^, with a cut off at 25 seconds to avoid tissue damage. Each paw was tested 3 times, with 5 min of rest between testing, and results were averaged for each animal.

To test <u>cold sensitivity</u>, animals were placed in small plexiglass enclosures on top of a thin 2.5 mm thick glass plate, and powdered dry ice packed into a 10 mL syringe with the top cut off was pressed against the glass beneath the plantar surface of the hindpaw^25^. Withdrawal latencies were recorded 3 times for each paw, with 5 min of break in between testing, and results were averaged for each animal. The maximum time allowed for withdrawal was 20 sec to avoid potential tissue damage.

<u>Spontaneous behaviors</u> were recorded in response to intraplantar Yoda-1 injections (10µM, 100µM and 1mM). Yoda1 or vehicle was injected into the plantar surface of the hindpaw and behaviors were recorded for 10 min following the injection. Videos were analyzed offline by an experimenter blinded to both genotype and treatment. Behaviors exhibited by the animals included biting and licking of the hindpaw.

### H&E staining

A hematoxylin and eosin stain was performed to assess the general morphology of the glabrous skin of Piezo1cKO and wildtype littermate controls. Glabrous skin of Piezo1cKO and wildtype animals was dissected and the tissue was fixed in 4% formaldehyde. The skin was processed, embedded in paraffin, sectioned into 4 μm sections and dried at RT until subsequent staining at the MCW Histology Core. Rehydrated sections were stained in hematoxylin for 3 min, washed in Richard-Allan Scientific™ Signature Series Clarifier™ 1,2 (for 45 sec, dipped for 30 sec in 0.1% ammonia water (bluing agent), stained in eosin for 30 sec, washed four times using 100% EtOH and lastly rinsed in Xylene. Slides were scanned using a Hamamatsu Nanozoomer HT slide scanner (Hamamatsu Photonics, K.K., Hamamatsu City, Japan) and images were assessed using NDP.View 2 software (Hamamatsu Photonics).

### Patch Clamp Recordings

On the third day of culture, keratinocyte recordings were made. Keratinocytes were superfused continuously with RT extracellular normal HEPES solution containing (in mM): 140 NaCl, 5 KCl, 2 CaCl2,1 MgCl2, 10 HEPES, and 10 glucose, pH 7.4 ± 0.05, and 310 ± 3 mOsm, and viewed on a Nikon Eclipse TE200 inverted microscope. Keratinocytes were patch clamped in both voltage clamp and current clamp mode (holding voltage -40 mV) with a borosilicate glass pipette (Sutter Instrument Company, Novato, CA) filled with intracellular normal HEPES solution containing (in mM): 135 KCl, 10 NaCl, 1 MgCl2, 1 EGTA, 0.2 NaGTP, 2.5 ATPNa2, and 10 HEPES, pH 7.20 ± 0.05, and 290 ± 3 mOsm. Cell capacitance and series resistance were maintained below 10 MΩ. Mechanical stimulation was elicited using a second borosilicate glass pipette that was driven by a piezo stack actuator (PA25, PiezoSystem Jena, Jena, Germany) at a speed of 106.25 μm/ms. Keratinocytes were stimulated with increasing displacements of 1.7 μm/Volt for 200 ms, and 2 min rest was allowed between displacements to avoid sensitization/desensitization of the cell membrane. Data was recorded using PatchMaster via an EPC10 amplifier HEKA Electronics, Holliston, MA). If the inward current elicited by the mechanical stimulations remained below 20 pA for at least three mechanical stimulations, the cell was considered mechanically insensitive. Data were analyzed using FitMaster (HEKA Electronics).

### Ex vivo Teased Nerve Fiber Recordings

Tibial skin nerve recordings were performed as previously described^26,3^. Briefly, animals were anesthetized and sacrificed via cervical dislocation. The leg of the animal was shaved and the glabrous skin with the innervating tibial nerve was quickly removed and placed in a heated (32 +-0.5°C), oxygenated bath (pH 7.45 +-0.05) consisting of (in mM): 123 NaCl, 3.5 KCl, 2.0 CaCl2, 0.7 MgSO4, 1.7 NaH2PO4, 5.5 glucose, 7.5 sucrose 9.5 sodium gluconate and 10 HEPES. Small nerve bundles were placed on a recording electrode and a blunt glass probe was used to search for receptive fields of single afferent fibers. Fibers were characterized based on their shape and conduction velocities: C-fibers < 1.2 m/s; Aδ-fibers 1.2–10 m/s; and Aβ-fibers for conduction velocities over 10 m/s^27^. Only slowly adapting Aβ and Aδ fibers were collected for these experiments. Action potential thresholds were determined using a continuous force ramp from 0 to 100 mN. A custom designed feedback-controlled mechanical stimulator was used to stimulate the receptive fields with 2, 5, 10, 20, 40, 100 and 150 mN for 10 seconds. Sensitization was prevented by allowing 1 min breaks between mechanical stimulations. Data was recorded and analyzed with LabChart (ADInstruments; Colorado Springs, CO).

### Data Analysis

Histological comparisons were made using a two-way ANOVA. For calcium imaging data comparing two groups, the percentage of keratinocytes responding was compared via Chi square and post hoc Fisher’s Exact tests; response magnitude was analyzed using one-way ANOVA with Bonferroni adjustment. For behavior experiments, paw withdrawal thresholds and suprathreshold stimulus responses were compared between two groups using non-parametric Mann-Whitney U-tests. Types of responses to the paintbrush and needle stimulus were analyzed using Chi square test with Fisher’s exact for groups of two. Spontaneous behavior was assessed using a Kruskal Wallis test or two-way ANOVA with Tukey’s post-hoc test. Paw withdrawal latencies were compared between two groups using the Student’s (two-tailed) *t* test. Skin nerve recordings were analyzed using a repeated measures two-way ANOVA with Sidak post-hoc test. Skin nerve mechanical thresholds were analyzed using Student’s (two-tailed) *t* test. Rheobase values and patch clamp mechanical thresholds were analyzed using a Mann-Whitney U-test, and resting membrane potentials were analyzed using the Student’s (two-tailed) *t* test. Current densities were analyzed using a mixed-effects analysis with Bonferroni post-hoc correction.

For all behavior experiments “n” corresponds to the number of animals. For patch clamp studies, skin nerve recordings or calcium imaging experiments at least n = 3 animals were utilized for each group shown, and the n on the graph corresponds to the number of cells, fibers, or repetitions. Summarized data are reported as mean ± SEM. The number within the bars on the graph corresponds to the number of animals used. All data analyses were performed using Prism 7 software (GraphPad, La Jolla, CA), with an alpha value of 0.05 set *a priori*. \**P* < 0.05, \*\**P* < 0.01, \*\*\**P* < 0.001, \*\*\*\**P* < 0.0001, n.s. denotes a non-significant comparison.

## Acknowledgement

The authors would like to thank Michael Lawlor, MD for assessing gross morphological differences between skin samples as well as Sarah Langer for experimental assistance. The authors also thank the Medical College of Wisconsin Histology Core for tissue sectioning and staining and the Medical College of Wisconsin Imaging Core for slide scanning.

## Author Contributions

FM planned, performed, and analyzed behavioral and electrophysiology experiments and wrote and edited the manuscript. ARM wrote and edited the manuscript.

KES planned, performed, and analyzed calcium imaging experiments and edited the manuscript. ADM planned, performed, and analyzed electrophysiology experiments and edited the manuscript. CLS assisted in experimental planning and manuscript editing.

